# NF-κB dependent expression of A20 controls IKK repression of RIPK1 dependent cell death in activated T cells

**DOI:** 10.1101/2023.09.22.558996

**Authors:** Scott Layzell, Alessandro Barbarulo, Benedict Seddon

## Abstract

The Inhibitor of Kappa B Kinase (IKK) complex is a critical regulator of canonical NF-κB activation. More recently, RIPK1 has also been identified as a phosphorylation target of the IKK complex, resulting in repression of extrinsic cell death pathways. Our previous work shows that normal thymocyte development is exclusively reliant on repression of TNF triggered cell death pathways by IKK, and that NF-κB signalling is in fact redundant for development. The role of IKK signalling in activated T cells is unclear. To investigate this, we analysed activation of IKK2 deficient TCR transgenic T cells with cognate peptide. While early activation events were normal, proliferation of blasts was impaired. Surprisingly, cell cycle progression in IKK2 KO T cells was unperturbed. Instead, dividing cells were more sensitive to apoptosis triggered by extrinsic cell death pathways, since inhibition of RIPK1 kinase activity almost completely rescued cell survival. Transcriptomic analysis of activated IKK2 deficient T cells revealed defective expression of several NF-κB targets, including Tnfaip3, that encodes A20, a negative regulator of NF-κB in T cells. To test whether A20 expression was required to protect IKK2 deficient T cells from cell death, we generated mice with T cells lacking both A20 and IKK2. Conditional deletion of both *Ikk2* and *Tnfaip3* in T cells resulted in near complete ablation of peripheral naïve T cells, in contrast to mice lacking one or other gene. Strikingly, this phenotype was completely reversed by inhibition of RIPK1 kinase activity in vivo. Therefore, our data suggests that IKK signalling in T cells protects against RIPK1 dependent death, both by direct phosphorylation of RIPK1 and through NF-κB mediated induction of A20, that we identify for the first time as a modulator of RIPK1 function in T cells.

## Introduction

The NF-κB family of transcription factors play critical roles in controlling development and function of many cell types (Bonizzi and Karin, 2004). Canonical NF-κB signalling is mediated by hetero or homodimers of p50, RELA and cREL family members that are sequestered in the cytoplasm by inhibitory proteins, the Inhibitors of kappa B (IκB) family and the related protein NFKB1. The key regulator of NF-κB dimer release is the inhibitor of kappa-B kinase (IKK) complex, a trimeric complex of two kinases, IKK1 (IKKα) and IKK2 (IKKβ), and a third regulatory component, NEMO (IKKγ). IKK phosphorylates IκB proteins, targeting them for degradation by the proteasome and releasing NF-κB dimers to enter the nucleus. During immune responses by T cells, activation of NF-κB is a critical early event following T cell receptor antigen recognition (reviewed in (Gerondakis and Siebenlist, 2010)). In the absence of REL subunits, or upstream NF-κB activators, such as Tak1 or IKK complex, T cells fail to blast transform or enter cell cycle (Webb et al., 2019; Xing et al., 2016). Consequently, mice with T cell specific ablation of RelA and/or cRel have peripheral naive T cells, but lack effector or memory phenotype T cells (Webb et al., 2019; Zheng et al., 2003). Similarly, disruption of the Card11-Bcl10-Malt1 (CBM) complex that links TCR triggering to NF-kB activation also prevents T cell activation (Egawa et al., 2003; Hara et al., 2003; Ruefli-Brasse et al., 2003; Ruland et al., 2001; Ruland et al., 2003)(Jost et al., 2007). Blast transformation of activated T cells is mediated by induction of c-Myc, and NF-κB is essential for c-Myc expression in mouse T cells (Grumont et al., 2004; Hayden and Ghosh, 2011).

Because of the critical function in T cell priming, it is less clear how NF-κB signaling contributes to later stages of T cell activation and effector differentiation. Analysis of upstream regulators of NF-κB activation provide some indications. Limited redundancy between IKK1 and IKK2 permits activation of T cells lacking one or other subunit (Chen et al., 2015; Schmidt-Supprian et al., 2003). However, IKK complexes formed by homodimers of either IKK1 or IKK2 are hypomorphic, and mice with T cell specific ablation of individual subunits exhibit reduced effector/memory compartments, most notably in IKK2 deficient mice (Schmidt-Supprian et al., 2003). Conversely, loss of negative regulators of NF-κB activation in mouse T cells result in de-repression of NF-κB activation and corresponding perturbations to T cell compartments. A20 is a potent suppressor of NF-κB activation. A20 deficient T cells have enhanced NF-κB activation that promotes anti-tumour CD8^+^ T cell responses (Giordano et al., 2014) and development of intrathymic Treg (Fischer et al., 2017). In spite of the expanded regulatory T cell pool, de-repression of NF-κB activation in these mice also results in increased numbers of effector and memory T cells (Onizawa et al., 2015). Several mechanisms have been proposed for how A20 inhibits NF-κB activation and specific activities may be cell or receptor context dependent. The deubiquitinase activity of A20 is thought to impair recruitment and/or retention of IKK to the CBM complex, by removal of K63-linked ubiquitin chains from MALT1(Duwel et al., 2009). Other studies suggest A20 deubiquitinase function is not required for regulation of TCR induced NF-κB in Jurkats. Rather, ZnF4/ZnF7 domains required for binding to K63 and M1 chains surrounding CBM complex mediate the suppressive function of A20. However, in other cell types, A20 can block TNF activation of IKK by binding polyubiquitin chains on NEMO subunit of IKK and thereby preventing IKK activation by Tak1, a mechanism largely dependent on A20’s seventh zinc-finger motif (ZnF7) (Skaug et al., 2011). Other studies suggest A20 competes with IKK for linear M1 ubiquitin chain binding using the ZnF7 domain.

Enhancement of memory and/or effector T cell responses may be mediated by pro-survival functions of NF-κB signalling. Optimal IL-2 synthesis by activated T cells is NF-κB-dependent and promotes their survival (Sriskantharajah et al., 2009) and addition of exogenous IL-2 can rescue defective Rel^−/−^ T cell responses to CD3 crosslinking in vitro (Saibil et al., 2007). BclXL has also been suggested to be an important pro-survival target of NF-κB during activation. In both mouse and human activated T cells, the expression of a dominant negative IκB construct results in impaired BclXL expression(Khoshnan et al., 2000; Mora et al., 2003). However, the importance of this for survival is questioned by the observation that T cell-specific ablation of BclxL does not result in death of activated T cells (Zhang and He, 2005).

A further complication to studying NF-κB pathways in T cells comes from the recent recognition that the IKK complex serves functions other than inducing NF-κB. Ablation of the IKK complex, either by deletion of NEMO (Schmidt-Supprian et al., 2003), or combined loss of IKK1 and IKK2 subunits (Webb et al., 2016) results in a developmental arrest in single positive (SP) thymocytes at the immature HSA^hi^ stage. Similar developmental blocks are observed in mice lacking the upstream activator of IKK, TAK1 (Liu et al., 2006; Wan et al., 2006; Xing et al., 2016). In contrast, expression of NF-κB REL subunits is not required for thymic development and generation of mature peripheral T cells (Oh et al., 2017; Webb et al., 2019). An explanation for this apparent contradiction comes from two key observations. First, the trigger for NF-κB activation via IKK in developing thymocytes is not TCR but TNF. The CBM complex but not required for thymocyte development or selection signalling (Jost et al., 2007; Schmidt-Supprian et al., 2004). In contrast, blockade of TNF signalling rescues development of IKK1/2 deficient thymocytes (Webb et al., 2016). Second, recent studies reveal that the IKK complex has two functions during TNF signalling in T cells - activating NF-κB and directly repressing cell death by inhibiting the serine threonine kinase, RIPK1. Ligation of TNFR1 causes recruitment of TRADD, TRAF2, and the serine/threonine kinase RIPK1. The ubiquitin ligases TRAF2, cellular inhibitor of apoptosis proteins (cIAPs) and the linear ubiquitin chain assembly complex (LUBAC), add ubiquitin chain modifications to themselves and RIPK1, creating a scaffold that allows recruitment and activation of the TAB/TAK and IKK complexes that in turn activate NF-κB. This is termed complex I (reviewed in (Annibaldi and Meier, 2018; Vandenabeele et al., 2010)). A failure to maintain the stability of this complex results in the formation of cell death inducing complexes. In the presence of IAP inhibitors, IKK inhibitors or TAK1 inhibitors (Dondelinger et al., 2013; Dondelinger et al., 2015; Wang et al., 2008), a complex composed of TRADD, FADD, CASPASE 8 and RIPK1 forms that induces apoptosis, a function dependent upon RIPK1 kinase activity (Annibaldi and Meier, 2018; Dondelinger et al., 2016; Ting and Bertrand, 2016). Phosphorylation of RIPK1 by IKK blocks RIPK1 kinase activity and therefore its capacity to induce apoptosis (Dondelinger et al., 2015). In thymocytes, it is this function of IKK, and not NF-κB activation, that is critical for their survival and onward development and accounts for the phenotype observed in IKK deficiency (Webb et al., 2019). A20 also appears to mediate cross-talk between NF-κB and cell death signalling pathways. In MEFs, A20 protects cells from death by binding and stabilizing the linear (M1) ubiquitin network associated to Complex I. Deletion of A20 induces RIPK1 kinase-dependent and -independent apoptosis upon single TNF stimulation (Priem et al., 2019). In T cells, A20 is proposed to protect cells from necroptotic cell death by restricting ubiquitination of RIPK3 and formation of RIPK1-RIPK3 complexes by its deubiquitinating motif (Onizawa et al., 2015).

In the present study, we investigated the role of IKK and NF-κB signalling pathways during T cell activation by analysing responses by IKK2 deficient T cells, that exhibit impaired IKK activity. Our results reveal that IKK signalling is critical to protect blasts from cell death by both transcriptional and non-transcriptional mechanisms and identify the NF-κB regulator A20 as a key target of NF-κB signalling activation that specifically attenuates cell death processes.

## Results

### IKK2 deficient CD8 T cells exhibit impaired proliferative responses to antigen stimulation

To better understand the role of NF-κB signalling pathways during T cell activation, we analysed antigen specific activation of IKK2 deficient T cells. In the absence of IKK2 expression, the IKK complex formed by IKK1 subunits is hypomorphic, but still sufficient to trigger canonical NF-κB activation (Schmidt-Supprian et al., 2003). To analyse antigen specific responses in the absence of IKK2, we analysed the F5.*Rag1*^-/-^ huCD2^iCre^*Ikk2^fx^* strain (F5*Ikk2*ΔT^CD2^ hereon) of TCR transgenic mice whose CD8^+^ T cells are specific for NP peptide from influenza, but lack IKK2 expression (Mamalaki et al., 1993; Silva et al., 2014). We tested reactivity of control and IKK2 deficient F5 T cells in vitro, to titrations of specific peptide, and in vivo, following their adoptive transfer to WT hosts and intranasal challenge with influenza A viral (IAV). In vitro, reactivity of F5 T cells was profoundly defective in the absence of IKK2 expression. Fewer T cells were observed in division over a range of peptide doses, though those cells that were triggered to divide matched the burst of divisions observed in controls (Fig. 1A). In vivo, F5 *Ikk2*ΔT^CD2^ T cells were activated in response to IAV infection, since they assumed a CD44^hi^CD62L^lo^ effector memory phenotype, but failed to accumulate either in lymphoid tissues or at the site of infection in the lung (Fig. 1B). Together, these results suggest a profound defect in the ability of T cells to become activated and generate abundant effectors in the absence of IKK2.

**Figure 1.**
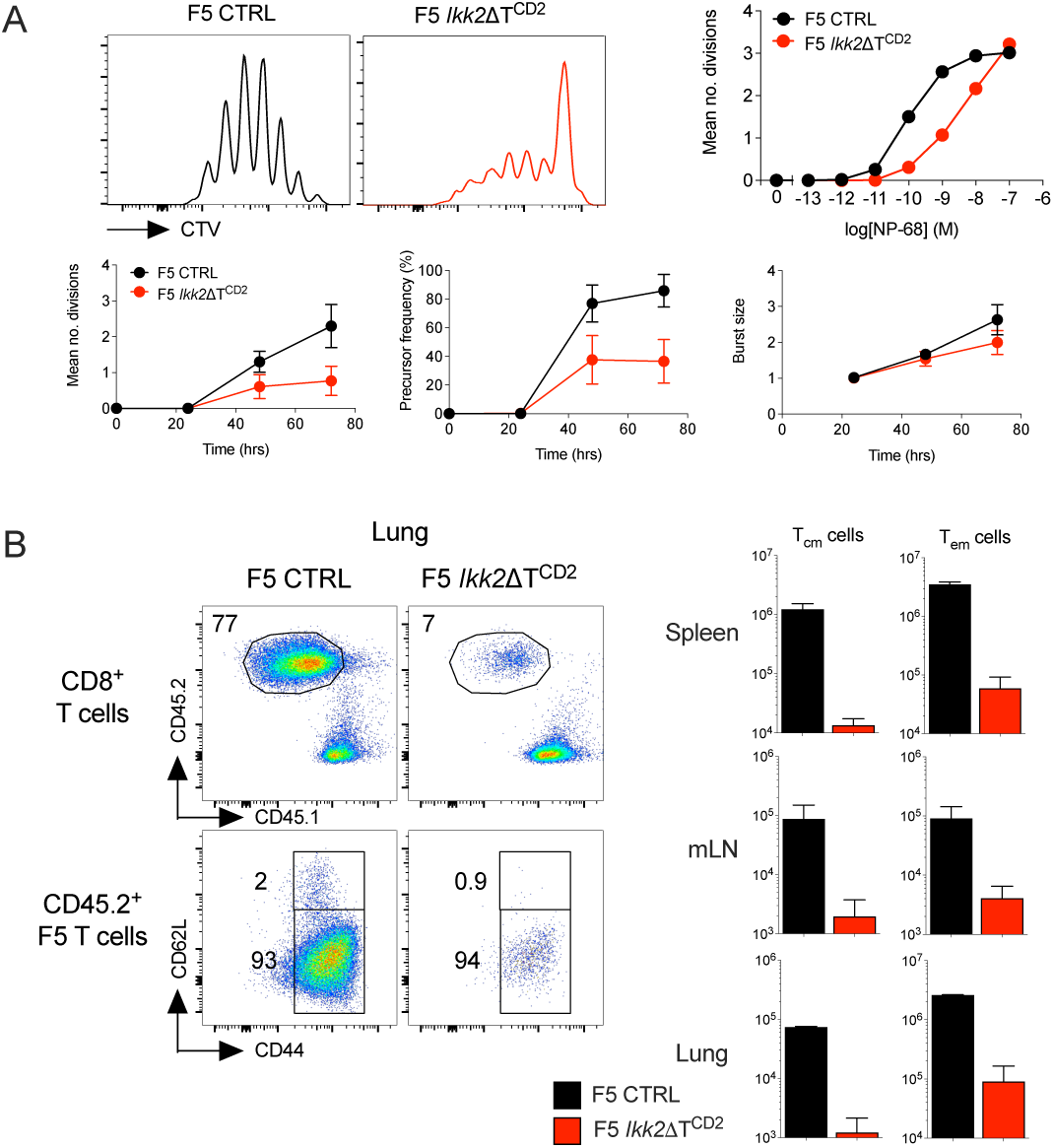
Defective activation of CD8^+^ F5 T cells in vitro and in vivo in the absence of IKK2. (A) CD8^+^ T cells were isolated from F5 *Ikk2*ΔT^CD2^ and Cre –ve controls, labelled with CTV cell dye and stimulated in vitro with a range of NP68 peptide concentrations. At different times, cells were recovered and CTV profiles analysed. Histograms show CTV labelling by T cells stimulated for 72h with NP68 at 10^-9^M. Line graphs show mean divisions at 72h over a range of peptide doses, and a time course of mean divisions, precursor of cells triggered into division and burst size of dividing cells, from cultures stimulated with NP68 10^-9^M. Data are representative of 5 independent experiments. (B) CD8^+^ T cells from F5 *Ikk2*ΔT^CD2^ and Cre –ve controls were transferred to CD45.1 WT hosts and challenged with IAV intranasally. 7 days later, lung, LN and spleen were recovered and CD45.2 donor T cell phenotype analysed. Density plots are of CD45.2 vs CD45.1 by CD8^+^TCR^hi^ cells, and CD62L vs CD44 expression by donor CD45.2 CD8^+^ TCR^hi^ T cells recovered from lung. Bar charts show total numbers of donor F5 T cells of CD62L^lo^CD44^hi^ TEM and CD62L^hi^CD44^hi^ TCM phenotype recovered from spleen, lymph node and lung of the same mice. Data are representative of two independent experiments.

### Normal triggering, blast transformation and cell cycle progression in the absence of IKK2

The reduced proliferation of F5 *Ikk2*ΔT^CD2^ T cells superficially resembled a failure of cells to be triggered following antigen recognition. To investigate this further, we examined early events of T cell activation in more detail. We first compared induction of CD69 and CD25 expression at 24h following peptide challenge, to determine whether sensitivity of TCR signalling was altered in the absence of IKK2. We observed similar induction of these early markers of triggering by F5.*Ikk2*ΔT^CD2^ and control T cells indicating that early activation events following TCR recognition were normal (Fig. 2A). We then examined blast transformation, analysing cell size increases and induction of cMyc at 24h, in response to peptide challenge. Both these measures were similar between F5 *Ikk2*ΔT^CD2^ T cells and controls (Fig. 2B), indicating that blast transformation was normal following antigen stimulation, in the absence of IKK2. We then examined cell cycle regulation in activated F5 T cells, comparing expression of key cell cycle regulators at 24h following activation, just prior to onset of cell division, and kinetics of the first cell division following activation. Expression of Cyclin D2 and D3, cyclin dependent kinase 6, Skp2 and E2f3 were all comparable between IKK2 deficient F5 T cells and controls. There was some evidence of a reduction in expression of cyclin dependent kinase inhibitor 1a in the absence of IKK2 (Fig. 2C). We compared the dynamics of cell cycle entry by measuring induction of Ki67 expression, that indicates entry of cells from G0 to G1 phase of cell cycle, DNA content to identify S phase and CTV cell dye labelling to identify cells that successfully complete first mitosis. In both control and IKK2 deficient T cell cultures, cells entered G1 between 18h and 30h, with a substantial fraction of these already in S phase by 30h. First division was completed by most cells by 38h. As indicated in earlier experiments, the fraction of F5 *Ikk2*ΔT^CD2^ T cells undertaking cell division was reduced. Nevertheless, we found no evidence that priming, blast transformation or cell cycle dynamics were defective in F5 *Ikk2*ΔT^CD2^ T cells. As such, the apparent proliferative defect must therefore be accounted for by defects in aspects of the T cell activation response other than cell cycle regulation.

**Figure 2.**
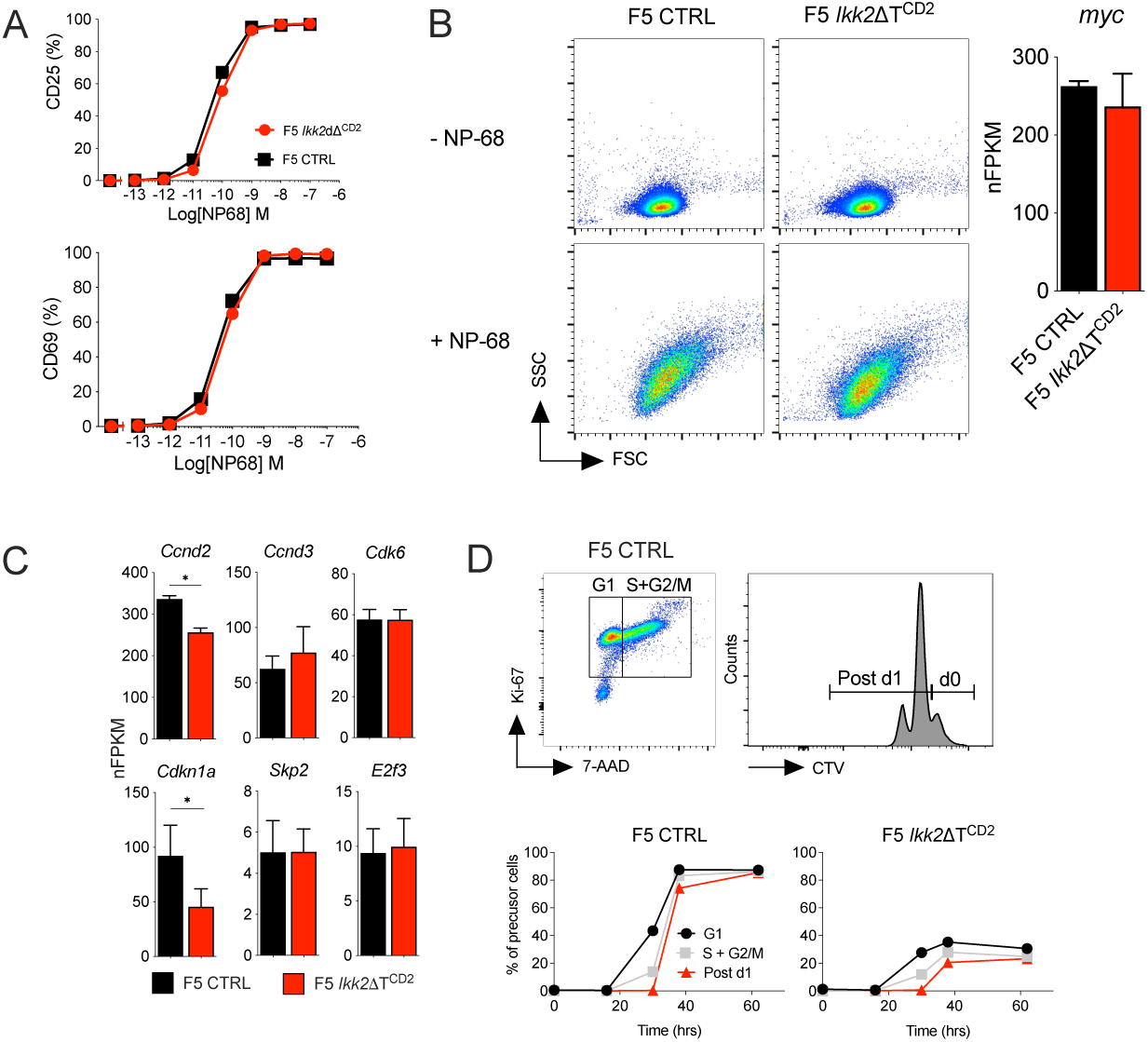
Normal priming and entry into cell cycle following activation of IKK2 deficient T cells. CD8^+^ T cells were isolated from F5 *Ikk2*ΔT^CD2^ and Cre –ve controls, labelled with CTV cell dye and stimulated in vitro with a range of NP68 peptide concentrations. (A) Line graphs show percent of cells expressing CD25 or CD69 at 24h of culture. (B) Density plots are of FSc vs SSc of T cells following 24h culture with NP68 at 10^-9^M. (B-C) Viable cells were recovered from cultures at 24h and RNA extracted. mRNA was analysed by bulk RNAseq. Bar chart shows Myc expression expressed as normalised FPKM (nFPKM) and cell cycle associated genes (C). (D) Cells stimulated with NP68 at 10^-9^M were harvested from cultures at different times and analysed for Ki67 expression and DNA content by 7-AAD staining. Density plot shows Ki67 vs 7-AAD staining by live gated CD8^+^ T cells in control cultures and gates used to define cells in G1 (KI67^hi^, diploid DNA content) and S+S2/M phase (Ki67^hi^ >diploid DNA content). Histogram shows CTV labelling at 38h and gates used to identify undivided cells and cells undergone 1 or more divisions. The line graphs are of precursor frequencies of undivided cells in G1 over time, vs cells in S+G2/M in undivided gate, vs cells that have completed their first division, changing with time for T cells from either Cre –ve controls or F5 *Ikk2*ΔT^CD2^ donors. Data are representative of four (B) or six (D) independent experiments.

### RIPK1 dependent cell death limits expansion of IKK2 deficient activated T cells

NF-κB has been implicated in controlling survival of T blasts by regulating intrinsic mitochondrial pathways of apoptosis through control of Bcl-XL expression (Khoshnan et al., 2000; Koenen et al., 2013; Mora et al., 2003). We therefore asked whether a cell death phenotype could account for the apparent proliferative defect of F5 *Ikk2*ΔT^CD2^ T cells. To do this, we assessed cell viability following T cell activation in vitro, by monitoring DNA content of cells. In cultures of control F5 T cells, a small fraction of dead cells was observed throughout the culture period. In contrast, we observed a progressive and substantial accumulation of dead cells in cultures of IKK2 deficient cells, peaking around 30h and significantly above levels observed in controls (Fig. 3A). Since NF-κB signalling may be perturbed in F5.*Ikk2*ΔT^CD2^ T cells, it was possible that alterations in expression of regulators of intrinsic pathways of apoptosis could be responsible for increased death of F5 *Ikk2*ΔT^CD2^ T cells. However, analysing expression of key repressors, activators and facilitators of intrinsic apoptotic pathways did not identify any gross defects in their gene expression (Fig. 3B).

**Figure 3.**
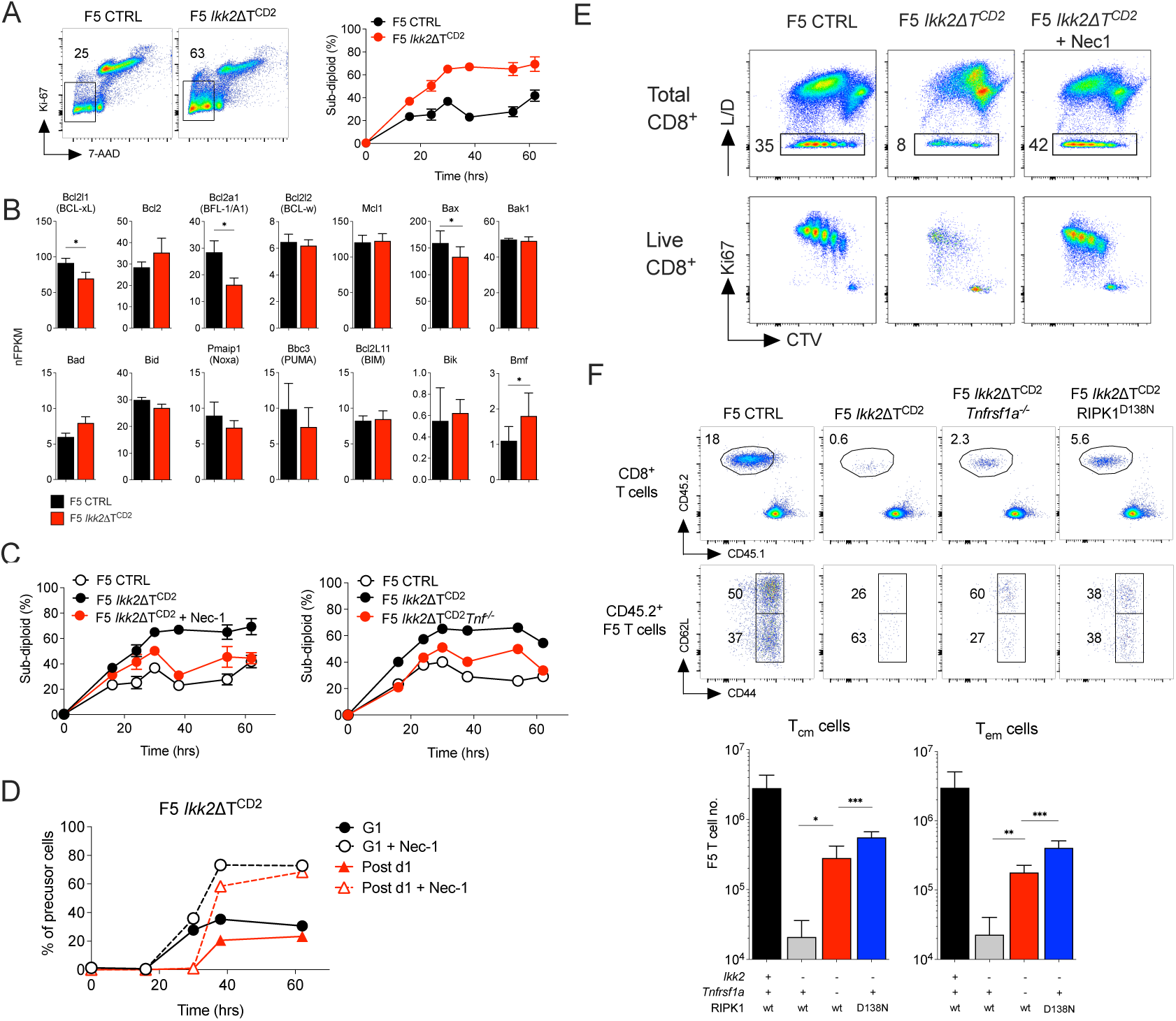
Induction of RIPK1 dependent cell death following activation of IKK2 deficient T cells. (A) CD8^+^ T cells were isolated from F5 *Ikk2*ΔT^CD2^ and Cre –ve controls, labelled with CTV cell dye and stimulated in vitro with 10^-9^M NP68 peptide. At different times during culture, T cells were recovered and stained for expression of Ki67 and DNA content by 7-AAD labelling. Density plots show Ki67 vs 7-AAD labelling at 38h of culture by total CD8^+^ cells. Gates indicate dead cells with sub-diploid DNA content. Line graphs are of % sub diploid cells from total CD8^+^ gate at different times during culture for T cells from the indicated mice. (B) Expression of genes that regulate intrinsic apoptotic cell death pathway, determined by RNAseq. Data are from the same experiment described in figure 2B-C. (C) Line graphs are of % sub diploid CD8^+^ cells at different times after culture of T cells from F5 *Ikk2*ΔT^CD2^ and Cre –ve controls, and cultures of F5 *Ikk2*ΔT^CD2^ T cells with addition of Necrostatin-1 (Nec1), and of similar cultures using T cells from F5 *Ikk2*ΔT^CD2^ and Cre –ve littermate controls and F5 *Ikk2*ΔT^CD2^ *Tnf*^-/-^ donors. (D) Entry of F5 T cells into cell cycle and completion of mitosis (Post d1) was analysed in cultures of T cells from F5 *Ikk2*ΔT^CD2^ donors in the presence and absence of Nec1. (E) Density plots are of CTV vs live/dead (L/D) on total CD8+ T cells, and CTV vs Ki67 by live CD8^+^ gated T cells in cultures at 72h. (F) F5 T cells from the indicated strains were transferred to groups of CD45.1 hosts and infected i.n with IAV. A d7, mice were culled and phenotype/ number of splenic T cells analysed. Density plots show donor CD45.2 vs host CD45.1 staining by live CD8^+^ T cells, and CD44 vs CD62L by donor CD45.2 F5 T cells from recipients of the indicated donor strains. Bar charts are of cell recovery of donor F5 T cells of Tcm and Tem phenotype from donor strains with the indicated combination of mutant alleles. Data are pool of three independent experiments (F).

Since we did not observe obvious defects in regulation of intrinsic apoptotic pathways, we next considered whether cell death was triggered by extrinsic pathways of apoptosis. TNF receptor superfamily members with death domains in the cytoplasmic tail of the receptor, are able to induce formation of a caspase 8 dependent death inducing complex. In the case of TNFR1, this complex also includes the serine/threonine kinase, RIPK1, whose kinase activity facilitates formation and activation of the Caspase 8 dependent cell death complex. To test whether death of F5 *Ikk2*ΔT^CD2^ T cells was triggered by extrinsic cell death pathways, we activated IKK2 deficient F5 T cells in the presence of RIPK1 kinase inhibitor, Necrostatin-1 (Nec1). Addition of Nec1 largely restored viability of activated F5 *Ikk2*ΔT^CD2^ T cells to levels observed in control cultures of IKK2^WT^ F5 T cells (Fig. 3C), implicating RIPK1 dependent extrinsic cell death pathways in the death of IKK2 deficient T cells. To ask whether TNF was a trigger of extrinsic cell death in F5.*Ikk2*ΔT^CD2^ T cells, we generated F5.*Ikk2*ΔT^CD2^*Tnf*^-/-^ mice. Loss of TNF expression by F5.*Ikk2*ΔT^CD2^*Tnf*^-/-^ T cells also restored T cell viability following activation in vitro (Fig. 3C). Blocking RIPK1 dependent extrinsic cell death pathways not only rescued F5 *Ikk2*ΔT^CD2^ T cell viability, but also restored cell proliferation by IKK2 deficient F5 T cells to levels comparable with control IKK2^WT^ F5 T cells, both in terms of fractions of cells successfully completing their first division (Fig. 3D) and the overall proliferative profiles later in cultures (Fig. 3E).

To confirm the relevance of these pathways in vivo, we tested whether loss of TNFR1 or RIPK1 kinase activity could rescue responses of F5.*Ikk2*ΔT^CD2^ T cells following IAV infection. We generated F5.*Ikk2*ΔT^CD2^*Ripk1^D138N^* mice expressing a kinase dead allele of RIPK1, and F5.*Ikk2*ΔT^CD2^*Tnfrsf1a^-/-^* mice that lacked expression of TNFR1. Measuring the response by different F5.*Ikk2*ΔT^CD2^ T cells following transfer into IAV challenged WT hosts revealed a substantial rescue of responding F5 T cell numbers 7d after infection (Fig. 3F). Taken together, these data reveal that, following their activation, F5 T cells become sensitised to TNF induced, RIPK1 dependent cell death in the absence of IKK2 expression.

### Reduced *Tnfaip3* expression by F5 *Ikk2*ΔT^CD2^ T cells following activation

The susceptibility of activated F5 *Ikk2*ΔT^CD2^ T cells to TNF induced RIPK1 dependent cell death was not predicted from earlier studies. The complete loss of IKK function resulting from ablation of IKK1/2 renders thymocytes and mature T cells highly susceptible to TNF induced cell death (Blanchett et al., 2022; Carty et al., 2022; Webb et al., 2016). However, expression of either IKK1 or IKK2 alone is sufficient to repress RIPK1 kinase activity and protect T cells. Accordingly, thymic development and peripheral naive T cell compartments are largely normal in mice whose T cells lack expression of only IKK1 or IKK2, in contrast to IKK1/2 deficient mice (Webb et al., 2019). One way to reconcile the present observations in activated F5 T cells with earlier studies would be if IKK signalling in these TCR transgenic cells were for some reason different to that of CD8^+^ T cells in WT polyclonal mice, in such a manner that IKK1 alone is not able to repress RIPK1 triggered cell death in F5 T cells. To test this, we cultured naive F5 T cells with increasing doses of TNF to see whether cells were susceptible to TNF induced cell death. Neither control nor F5.*Ikk2*ΔT^CD2^ T cells were induced to undergo cell death even at supra-physiological concentrations of TNF, suggesting that IKK1 alone is sufficient to protect F5 T cells (Fig. 4A). To further confirm that IKK signalling was required to protect F5 T cells from RIPK1 triggered cell death, we also tested whether IKK inhibitor could sensitise F5 T cells to TNF induced death. Addition of IKK inhibitor did indeed sensitise F5 T cells to TNF induced cell death even at low doses of TNF (Fig. 4A). Addition of Nec1 to cultures blocked cell death, confirming death was RIPK1 dependent. Taken together, these results suggest that F5 *Ikk2*ΔT^CD2^ T cells become sensitive to TNF induced cell death following their activation, since naive F5 *Ikk2*ΔT^CD2^ T cells were resistant to TNF induced cell death. Development of sensitivity to TNF induced cell death must therefore result from changes in regulation of death signalling during T cell activation.

**Figure 4.**
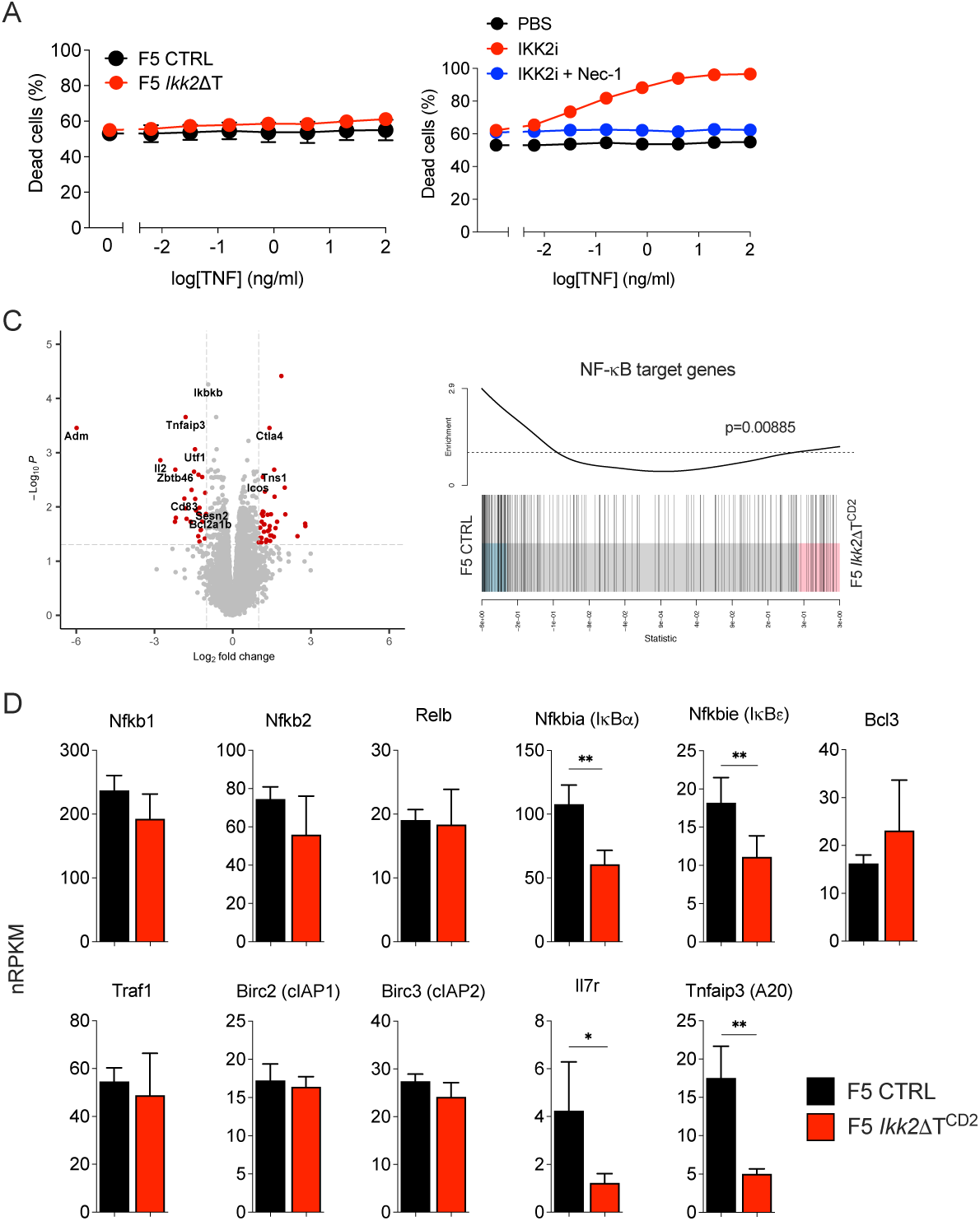
Impaired expression of *Tnfaip3* by IKK2 deficient T cells following antigen stimulation. CD8^+^ T cells were isolated from F5 *Ikk2*ΔT^CD2^ and Cre –ve controls were cultured overnight with TNF and viability determined by viability dye staining. Control cells were additionally cultured with IKK2 inhibitor (IKK2i, 10µM) and Nec1 (10µM). (B) Volcano plot comparing gene expression of IKK2 deficient F5 T cells vs control F5 T cells 24h after activation. (C) Gene set enrichment analysis comparing NF-κB regulated gene expression between activated control and IKK2 deficient F5 T cells. (D) Bar charts show expression of NF-κB gene targets specifically validated in T cells. Data are representative of three or more independent experiments (A).

Our earlier studies show that TNFRSF induced NF-κB activation is reduced in F5 *Ikk2*ΔT^CD2^ T cells that only express IKK1(Silva et al., 2014). Therefore, we hypothesised that activated F5.*Ikk2*ΔT^CD2^ T cells fail to induce expression of NF-κB target gene(s) necessary to protect them from extrinsic cell death. To identify potential candidate genes we performed differential gene expression analysis of control and F5.*Ikk2*ΔT^CD2^ T cells 24h after activation (Fig. 4C). Gene enrichment analysis confirmed the presence of an NF-κB target gene signature in control T cells that was reduced in F5 *Ikk2*ΔT^CD2^ T cells. However, a focused analysis of validated NF-κB gene targets specifically identified in IKK1/2 deficient thymocytes (Webb et al., 2019), revealed normal expression of many target genes by activated F5.*Ikk2*ΔT^CD2^ T cells. In thymocytes, expression of *Nfkb2, Relb, Bcl3, Traf1, Birc2* and *Birc3* is NF-κB dependent, but were all normally expressed in activated F5.*Ikk2*ΔT^CD2^ T cells (Fig. 4D). This suggests that IKK1 mediated activation of NF-κB was in many cases sufficient for normal NF-κB dependent gene expression. There were, however, a number of defined NF-κB target genes in T cells whose expression was impaired in F5 *Ikk2*ΔT^CD2^ T cells, such as *Nfkbia, Il2, Il7r* and *Tnfaip3,* which encodes A20 protein.

### Loss of A20 expression sensitises T cells to IKK dependent TNF induced cell death

The defect in *Tnfaip3* expression was notable, since A20 protein, encoded by *Tnfaip3*, can inhibit NF-κB activation but is also implicated in regulating apoptotic and necroptotic cell death signalling in MEFs and T cells respectively. To investigate which of these functions was relevant in activated T cells in our studies, we first analysed T cells from *Tnfaip3^flox^ Cd4^Cre^* (*Tnfaip3ΔT^CD4^*) mice whose T cells lack A20 expression. Consistent with earlier studies, we found that, while total T cell numbers were largely normal in the absence of A20, there were significant increases in numbers of CD4 memory and Foxp3^+^ regulatory T cell populations (Fig. 5A). Treg numbers were also increased in the thymus, suggesting enhanced production in the absence of A20. TCR triggered NF-κB signalling is implicated in generation of both regulatory T cells and CD4 memory populations, so the observed increases were entirely consistent with a dis-inhibition in NF-κB signalling in the absence of A20. We have previously shown that NF-κB activation by TNFSRF members is important for induction of *Il7r* expression and naive T cell survival (Silva et al., 2014). Analysing naive T cell numbers and IL7R protein did not reveal any evidence of a comparable de-repression of NF-κB signalling downstream of TNFSRF in *Tnfaip3ΔT^CD4^* mice.

**Figure 5.**
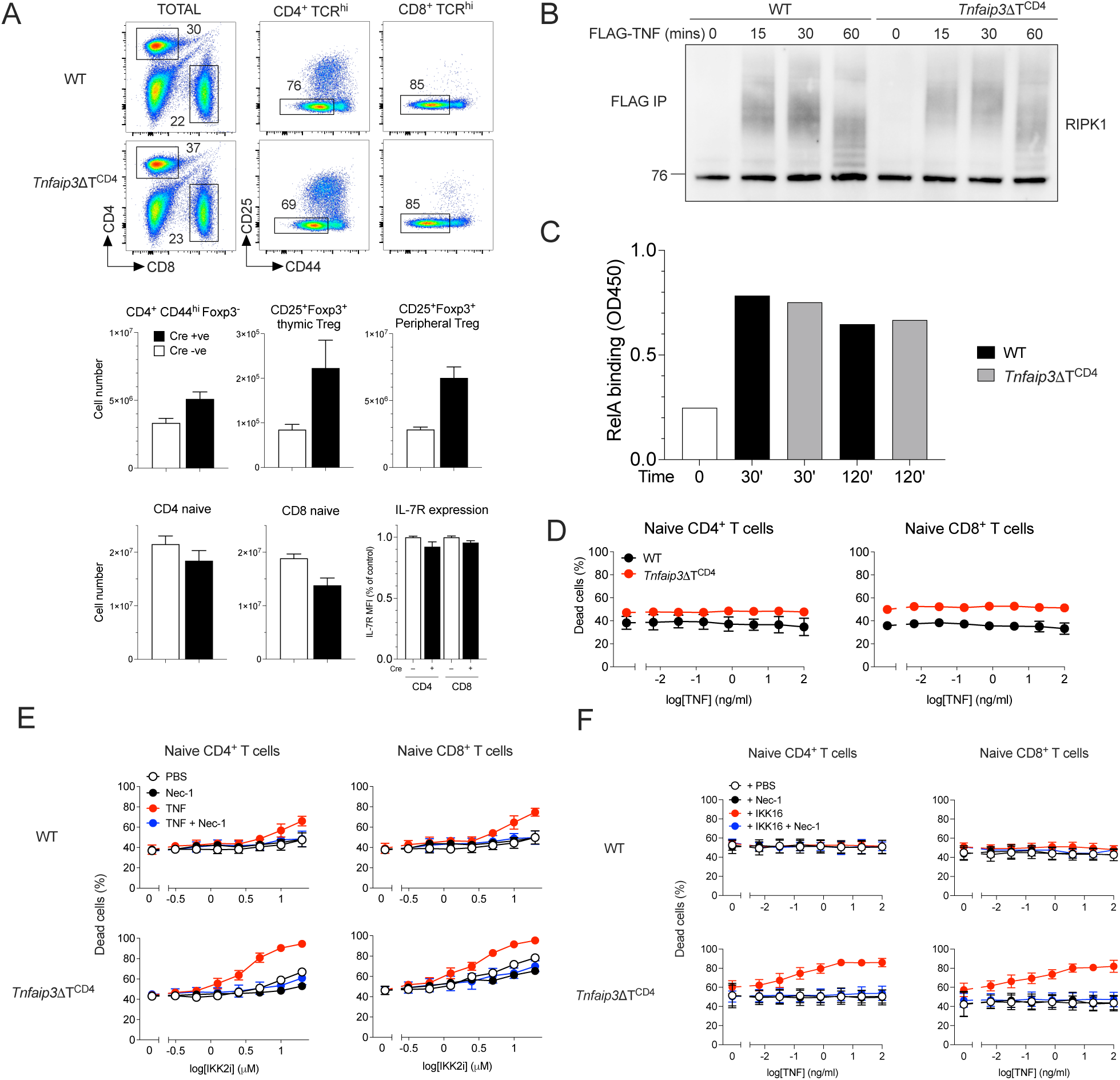
Normal NF-κB activation but defective regulation of extrinsic cell death pathways in *Tnfaip3* deficient T cells. (A) Lymph nodes and spleen were recovered from *Tnfaip3*ΔT^CD4^ and Cre –ve litter mates aged between 8 and 12 weeks, and T cell phenotype and number determined by flow cytometry. Density plots are of CD4 vs CD8 expression by total live lymph node cells, CD25 vs CD44 by the indicated subset. Bar charts summarise total numbers of TCR^hi^CD4^+^ memory phenotype (CD4^+^CD44^hi^Foxp3^−^), Treg (CD25^+^Foxp3^+^ Treg), CD4^+^TCR^hi^CD44^lo^ naive T cells (CD4 naive) and CD8^+^TCR^hi^CD44^lo^ naive T cells (CD8 naive) recovered from total lymph nodes and spleen combined. Total numbers of CD25^+^Foxp3^+^ thymic Treg are from thymus. Bar chart of IL-7R expression by naive CD4 and naive CD8 T cells is expressed as fraction of MFI normalised to MFI of corresponding subset from Cre–ve controls. (B) T cells from *Tnfaip3*ΔT^CD4^ and Cre –ve litter mates were stimulated with FLAG-TNF for different times, lysed and immunoprecipitated with anti-FLAG-sepharose beads and analysed by immunoblotting for RIPK1. (C) T cells from *Tnfaip3*ΔT^CD4^ and Cre –ve litter mates were stimulated with TNF for 30’ or 120’, nuclear extracts prepared and RELA levels determined by ELISA. (D-F) Lymph node cells from *Tnfaip3*ΔT^CD4^ and Cre –ve litter mates were cultured overnight with titrations of TNF, IKK2 inhibitor (IKK2i), in the presence or absence of fixed concentrations of TNF (20ng/ml), Nec1 (10µM), pan IKK inhibitor (IKK16, 0.25µM). Line graphs show fraction of dead cells amongst TCR^hi^ naive CD4^+^CD44^lo^ and TCR^hi^ naive CD8^+^CD44^lo^ T cells from *Tnfaip3*ΔT^CD4^ and Cre –ve litter mates from cultures titrating TNF (D), in cultures titrating IKK2i in the presence or absence of Nec1 and/or TNF (E) and in cultures titrating TNF with addition of Nec1 and or IKK16 (F). Data are representative (A-C) or show average of three or more independent experiments. Error bars show SEM.

In HEK293 cells and MEFs, the deubiquitinase function of A20 has been implicated in destabilising TNFR complex I and triggering cell death (Wertz et al., 2004), while other studies in MEFs show A20 binds and stabilizes the linear (M1) ubiquitin network associated to complex I, thereby inhibiting complex II formation and TNF induced cell death(Draber et al., 2015; Priem et al., 2019). Since TNF induced cell death of IKK2 deficient effectors was RIPK1 dependent, we first analysed RIPK1 recruitment and ubiquitination in TNFR complex I in A20 deficient T cells. Total thymocytes from control and *Tnfaip3ΔT^CD4^* mice were stimulated with FLAG-TNF and complex I immunopreciptated at different timepoints following stimulation. Blotting for RIPK1 revealed the dynamics of RIPK1 ubiquitination over time. Analysing WT thymocytes revealed heavy ubiquitination of RIPK1 that gradually declined at later timepoints (Fig. 5B). A20 deficient thymocytes exhibited similar levels and extent of ubiquitination of RIPK1 following stimulation as WT. Analysing NF-kB nuclear translocation following TNF stimulation revealed similar levels and kinetics of NF-kB activation in A20 deficient T cells as controls (Fig. 5C). We also analysed viability of A20 deficient T cells following TNF stimulation. In contrast to MEFs, A20 deficient T cells remained resistant to TNF induced cell death (Fig. 5D). Since IKK activity is essential to repress RIPK1 mediated cell death in T cells, we measured survival of A20 deficient T cells with TNF when IKK is inhibited. Titrating IKK2 inhibitor revealed that A20 deficient T cells were highly sensitive to IKK inhibition compared with controls (Fig. 5E), and were similarly highly sensitive to low levels of TNF with suboptimal IKK inhibition (Fig. 5F). Cell death was blocked by Nec1, confirming RIPK1 dependence. Taken together, these data suggested that in the absence of A20 expression, the capacity of IKK to repress RIPK1 induced cell death is greatly impaired, such that even partial loss of IKK function and very low levels of TNF are sufficient to induce cell death.

### A20 is critical for survival of IKK2 deficient T cells in vivo

Our analysis of A20 deficient T cells suggested that IKK dependent repression of RIPK1 is highly dependent upon A20 for optimal activity. To test for evidence of epistasis between *Tnfaip3* and *Ikbk2*, we generated *Ikbkb2^flox^ Tnfaip3^flox^ Cd4^Cre^* (*Tnfaip3.Ikk2ΔT^CD4^*) mice whose T cells lack expression of both IKK2 and A20 protein. In mice lacking one or other of A20 or IKK2, thymic development is normal and T cell compartments are largely normal in size, other than a modest reduction in the naive CD8 compartment in *Ikbkb2^flox^ Cd4^Cre^* mice (Fig. 6A-B) that has been previously reported (Silva et al., 2014). Analysing thymi of *Tnfaip3.Ikk2ΔT^CD4^* mice first, revealed normal representation of most major DP and SP subsets. However, a small but significant reduction in the most mature CD62L^hi^HSA^lo^ stage of CD8 T cell development was evident (Fig. 6A). In contrast, quantifying peripheral T cell compartments revealed a potent epistatic interaction between *Tnfaip3* and *Ikbk2*. As compared with controls or single knockout strains, ablation of both IKK2 and A20 resulted in profound lymphopenia in both CD4^+^ and CD8^+^ T cell subsets (Fig. 6B).

**Figure 6.**
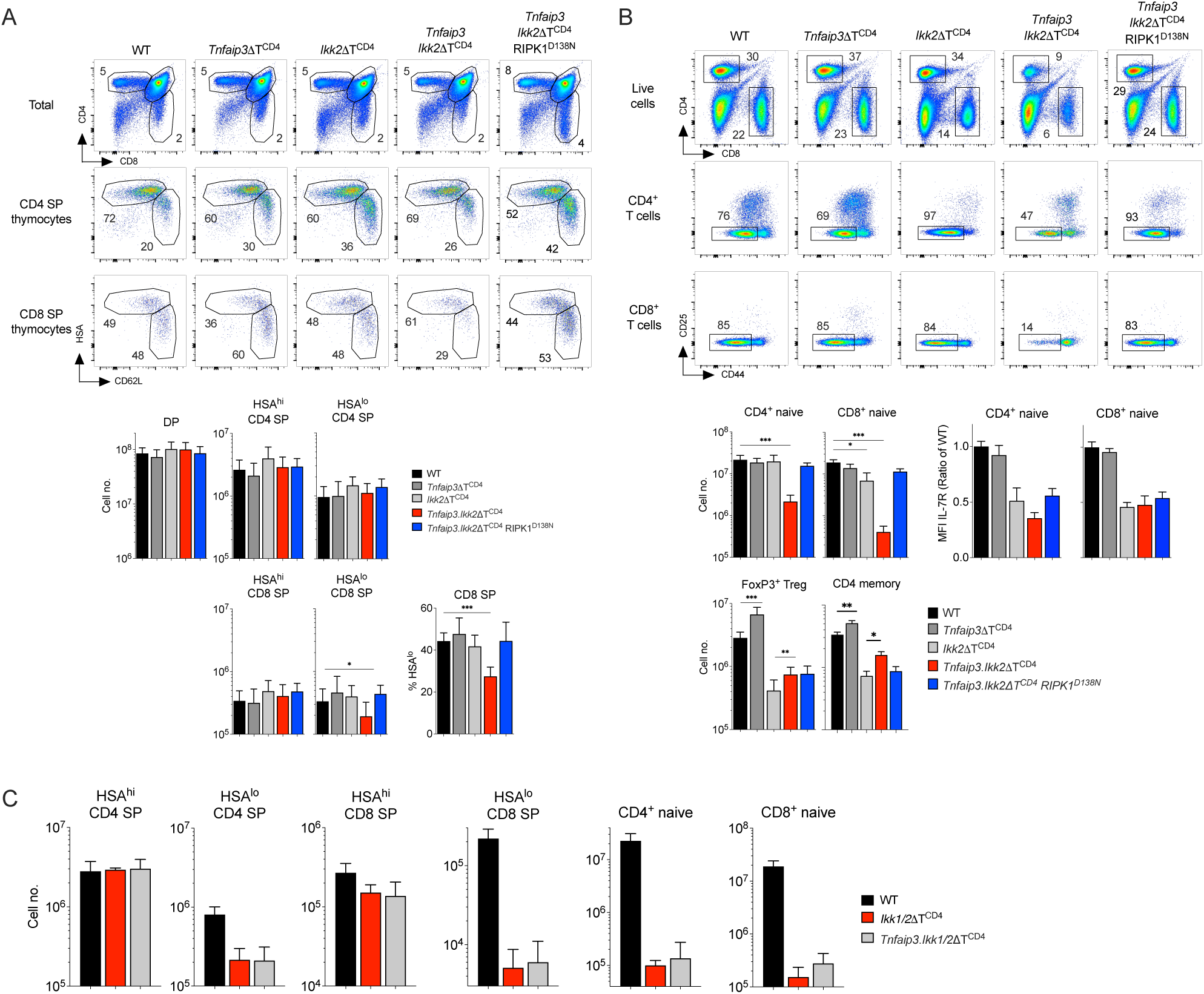
*Tnfaip3* expression by IKK2 deficient T cells is crucial for repression of RIPK1 induced cell death. Thymus, lymph nodes and spleens were recovered from *Tnfaip3*ΔT^CD4^, *Ikk2*ΔT^CD4^, *Tnfaip3.Ikk2*ΔT^CD4^,*Tnfaip3.Ikk2*ΔT^CD4^RIPK1^D138N^, *Ikk1.Ikk2*ΔT^CD4^ and *Tnfaip3. Ikk1.Ikk2*ΔT^CD4^ mice between 10-16 weeks of age. Composition and size of T cell compartments was determined by flow cytometry. (A) Density plots are of CD4 vs CD8 by total live thymocytes, HSA vs CD62L by CD4^+^CD8^−^TCR^hi^ (CD4 SP thymocytes) and CD4^−^CD8^+^TCR^hi^ (CD8 SP thymocytes) from the indicated strains. Bar charts show total numbers of the indicated subset in different strains, and for CD8 SP thymocytes, the frequency of mature HSA^lo^ thymocytes amongst total CD8 SP thymocytes. (B) Density plots are of CD4 vs CD8 by total live lymph node cells, CD25 vs CD44 by gated CD4^+^TCR^hi^ (CD4^+^ T cells) and CD8^+^TCR^hi^ (CD8^+^ T cells) cells from lymph nodes of the indicated strains. Bar charts show total numbers recovered from lymph nodes and spleen combined, of the indicated subsets in different strains, and IL-7R expression level, expressed as a fraction of the same subset from Cre–ve controls analysed on the same day, for CD4^+^CD44^lo^ and CD8^+^CD44^lo^ naive T cells recovered from lymph nodes. (C) Bar charts show cell numbers of thymic subsets of CD4 and CD8 SPs, and peripheral CD4^+^CD44^lo^ (CD4^+^ naive) and CD8^+^CD44^lo^ (CD8^+^ naive) T cells recovered from *Ikk1.Ikk2*ΔT^CD4,^ *Tnfaip3. Ikk1.Ikk2*ΔT^CD4^ and Cre–ve littermates mice. Numbers of mice are indicated in the legends and are pooled from 4 or more independent analyses.

We next asked whether the T lymphopenia in *Tnfaip3.Ikk2ΔT^CD4^* mice was the result of a RIPK1 induced cell death process, by testing if kinase dead RIPK1 would block cell death and reverse lymphopenia. To do this we generated and analysed *Tnfaip3.Ikk2ΔT^CD4^ Ripk1^D138N^* mice. In vivo inactivation of RIPK1 kinase activity resulted in a near complete reversal of T lymphopenia in *Tnfaip3.Ikk2ΔT^CD4^* mice. Numbers and representation of mature CD62L^hi^HSA^lo^ CD8 SP thymocytes were restored by introduction of RIPK1^D138N^ mutation (Fig. 6A). Similarly, RIPK1^D138N^ mutation restored numbers of naive CD4^+^ and naive CD8^+^ T cells in *Tnfaip3.Ikk2ΔT^CD4^* mice to near normal levels, confirming the dominant role of extrinsic cell death processes in the lymphopenia of *Tnfaip3.Ikk2ΔT^CD4^* mice.

Taken together, these data suggest a powerful synergy between IKK2 and A20 in regulating resistance to cell death not evident in single deficient strains. Since A20 deficient T cells were more sensitive to TNF induced cell death in vitro (Fig. 5), we hypothesised that A20 was required to facilitate efficient repression of RIPK1 by IKK. If so, then impact of A20 ablation on cell death should be dependent on IKK activity and in the complete absence of IKK, further A20 ablation would have no impact. To test this, we generated *Chuk^flox^ Ikbkb^flox^ Tnfaip3^flox^ CD4^Cre^* (*Ikk1.Ikk2.Tnfaip3*ΔT^CD4^) mice that lack expression of both the IKK complex and A20. If A20 is in fact acting independently of IKK, we would expect an additive effect of A20 deficiency upon IKK deficiency, as observed when comparing *Ikk2ΔT^CD4^* and *Tnfaip3.Ikk2ΔT^CD4^* mice. In mice lacking IKK expression, thymic development was profoundly blocked at the mature SP stage, and mice had reduced numbers peripheral naive T cells (Fig. 6C), as previously reported(Schmidt-Supprian et al., 2003; Webb et al., 2016). Analysing *Ikk1.Ikk2.Tnfaip3*ΔT^CD4^ mice that lack A20 and the IKK complex, revealed a phenotype indistinguishable from that of *Ikk1.Ikk2*ΔT^CD4^ mice. Additional loss of A20 did not further exacerbate the phenotype of *Ikk1.Ikk2*ΔT^CD4^ mice.

### Accelerated cell death of activated T cells lacking both IKK2 and A20

Following T cell activation, TCR induced TRAF activity targets A20 for degradation (Coornaert et al., 2008; Duwel et al., 2009). Our data suggested that IKK2 deficient T cells fail to successfully induce re-expression of A20, resulting in their sensitisation to RIPK1 dependent cell death. To validate this, we analysed survival of A20/IKK2 deficient T cells following activation to determine whether their behaviour phenocopied that of activated IKK2 deficient T cells. T cells were isolated from *Ikk2ΔT^CD4^*, *Tnfaip3.Ikk2ΔT^CD4^* and *Tnfaip3.Ikk2ΔT^CD4^ Ripk1^D138N^* mice, labelled with CTV cell dye and activated by plate bound CD3 and CD28. Analysing proliferation and cell death revealed the anticipated impairment of T cell viability amongst both CD8^+^ and CD4^+^ T cells from *Ikk2ΔT^CD4^* mice. Consistent with experiments analysing peptide specific activation of IKK2 deficient F5 T cells, we did not observe substantial RIPK1 dependent cell death at 24h. However, by 48h, cultures of IKK2 deficient CD4^+^ and CD8^+^ T cells exhibited low levels of viability that could be restored by inhibiting RIPK1 with Nec1 (Fig. 7). By comparison, T cells from *Tnfaip3.Ikk2*ΔT^CD4^ mice exhibit high levels of cell death even by 24h, that were blocked by introduction of kinase dead RIPK1^D138N^. By 48h, levels of death in cultures of T cells from *Tnfaip3.Ikk2*ΔT^CD4^ mice resembled those from *Ikk2ΔT^CD4^* mice.

**Figure 7.**
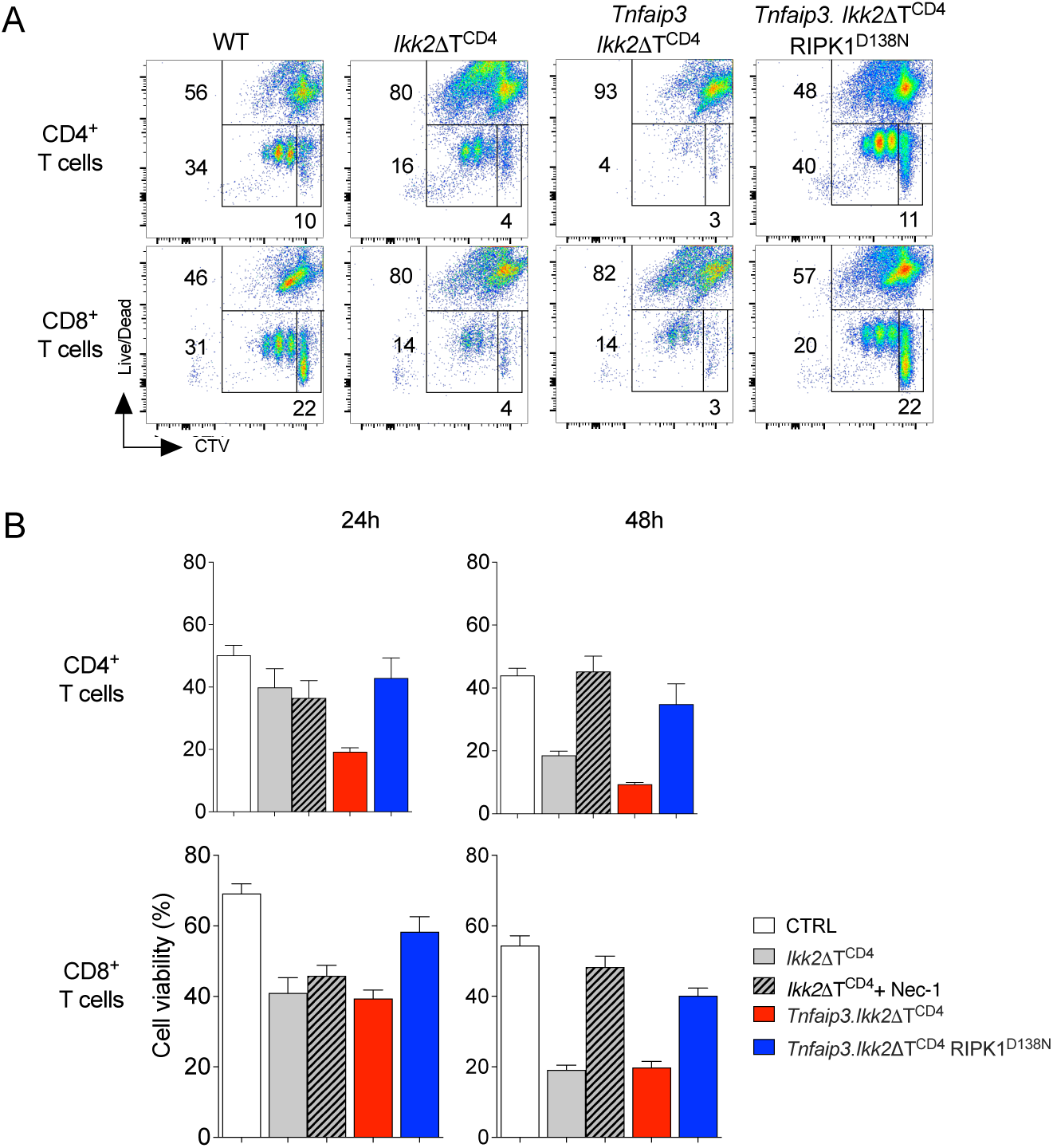
Accelerated cell death of activated T cells lacking IKK2 and A20. T cells from lymph nodes of Ikk2ΔTCD4, Tnfaip3.Ikk2ΔTCD4, Tnfaip3.Ikk2ΔTCD4RIPK1D138N strains, and Cre–ve litter mates were labelled with CTV and activated in vitro with plate bound anti-CD3 and CD28. Viability of CD4^+^ and CD8^+^ T cells was assessed at 24h and 48h of culture by flow cytometry. T cells from Ikk2ΔTCD4 donors were additionally cultured with and without Nec1 (10µM). (A) Density plots are of Live/dead dye vs CTV dilution profile of total CD4^+^ or CD8^+^ T cells from the indicated strains. (B) Bar charts show % live cells in cultures of the indicated T cell subset from the indicated strain and condition at either 24h or 48h post activation. Data are representative (A) or mean +/- SEM (B) of four independent experiments.

## Discussion

In this study, we investigated the roles of NF-κB and cell death signalling following T cell activation, in the development of effector responses specifically in the period that follows initial priming, which is strictly NF-κB dependent. IKK2 deficient T cells provide a sensitised model of hypomorphic IKK signalling during T cell activation, in which reduced IKK activity is sufficient for initial priming, but reveals adaptive regulation of extrinsic cell death pathways, mediated by NF-κB dependent control of the gene for A20, *Tnfaip3*. Our results suggest a new function for A20 in the regulation of extrinsic cell death pathways in T cells, by facilitating optimal repression of RIPK1 mediated cell death by IKK complex and that is controlled by NF-κB dependent expression of *Tnfaip3* during activation.

Our previous work showed that IKK signalling is essential for survival of mature thymocytes, by phosphorylating and thereby repressing the kinase activity of RIPK1. In the absence of IKK activity, thymocytes die by TNF induced RIPK1 triggered apoptosis(Webb et al., 2019). In thymocytes and naive T cells, it appears a relatively low level of IKK activity is sufficient for this survival function, since hypomorphic IKK complexes that form in the absence of either IKK1 or IKK2 expression, are sufficient to maintain control RIPK1 dependent cell death. Mice with T cell specific deletion of *Ikk1* or *Ikk2* exhibit normal thymocyte and peripheral T cell survival(Webb et al., 2019). In confirmation, we found here that IKK2 deficient naive F5 T cells were resistant to TNF induced cell death even at supra-physiological levels of TNF. On this basis, the observed susceptibility of IKK2 deficient T cells to RIPK1 dependent TNF induced cell death following activation was not predicted, but suggested a re-configuration of extrinsic cell death pathways, as a consequence of T cell activation, that heightened sensitivity to RIPK1 dependent cell death.

NF-κB has been implicated in regulating survival of many cell types, including T cells, by tuning expression of Bcl2 family members that control the intrinsic mitochondrial pathway of cell death. This and previous studies of IKK2 deficient T cells (Silva et al., 2014) did not find evidence of perturbed mitochondrial cell death regulation. We observed only a modest reduction of A1 expression in activated IKK2 deficient T cells. Our results do not necessarily exclude a role for NF-κB regulation of intrinsic cell death pathways in T cells and may rather reflect variation in the thresholds of NF-κB activity required to regulate expression of different target genes. Our previous studies of REL and IKK deficient T cells identified a number of genes whose expression was NF-κB dependent in T cells, with clear reductions of expression observed in vivo(Webb et al., 2019; Webb et al., 2016). While it is clear that activation of NF-κB is suboptimal in either IKK1 or IKK2 deficient T cells (Schmidt-Supprian et al., 2003; Webb et al., 2016), we found that expression of many NF-κB gene targets, including *Myc* that is required for blast transformation, and *CIap1/2*, required for survival, were normal in IKK2 deficient T cells following activation. In contrast, *Il7r*, and negative regulators of NF-κB signalling, *Ikbia*, *Ikbie* and most notably *Tnfaip3*, were reduced. Our results reveal a hierarchy of NF-κB target gene regulation in T cells, with a subset of target genes exhibiting a higher threshold of NF-κB transcriptional activity required for optimal expression.

Following activation, IKK2 deficient T cells appeared to lose control of RIPK1, allowing RIPK1 to trigger cell death. We ascribe a transcription mechanism to this change in death signaling, resulting from a failure of activated IKK2 deficient T cells to restore NF-κB dependent expression of *Tnfaip3*. Analysing T cells lacking A20 revealed a novel function of this key regulator in controlling extrinsic cell death pathways, regulating the IKK-RIPK1 signaling axis. A20 is implicated in regulating both NF-κB activity and cell death and so a loss of *Tnfaip3* expression following activation of IKK2 deficient T cells could have had multiple consequences. In T cells, there is clear evidence from A20 deficient mice for a loss of negative feedback regulation of NF-κB activation in vivo, since Treg and CD4^+^ memory phenotype cells are increased in number and there are several reports of enhance effector function by A20 deficient T cells in vivo(Giordano et al., 2014; Just et al., 2016). Notably, both these subsets and functions rely upon TCR triggered NF-κB(Ruland et al., 2001). It appears that in T cells, the negative feedback function of A20 is restricted to TCR induced NF-κB. We did not observe any evidence that TNF induced NF-κB activity or expression of gene targets such as IL7R, was increased in T cells in the absence of A20. In terms of cell death control, A20 has been implicated in control cell death pathways in MEFs mediated by binding and stabilizing of the M1 linear ubiquitin network associated with complex I. Deletion of A20 appears to be sufficient to destabilise the M1 ubiquitin network resulting in both RIPK1 kinase-dependent and -independent apoptosis upon single TNF stimulation(Priem et al., 2019). In T cells, A20 deletion alone does not appear to be sufficient destabilise complex I formation to the point of death, since *Tnfaip3ΔT^CD4^* mice had normal T cell numbers. However, stressing cell death signalling in T cells with suboptimal levels of IKK inhibitors did reveal an underlying fragility in complex I. In IKK2 deficient T cells, further loss of A20 in both resting naive T cells or in activated T cells was sufficient to render cells highly susceptible to TNF induced cell death. In contrast to MEFs, cell death was entirely RIPK1 dependent, and depends on IKK kinase activity. A20 ablation had no additional impact on cell viability in the complete absence of IKK function. The most parsimonious explanation of these data is that A20 expression is required for optimal interaction of IKK with RIPK1 to allow efficient phosphorylation and repression of RIPK1 kinase activity. Whether this function relies on M1 binding capacity of A20, as is the case in MEFs, remains to be determined.

Our results also shed light on the dynamics and triggers of NF-κB signalling during T cell activation. Initial TCR stimulation is required to activate T cells, and induction of NF-κB targets such as Myc are critical for early events such as blast transformation. However, our data also clearly demonstrate the activity of TNF later in the response, revealed by cell death events in the absence of IKK2 expression. However, in normal T cells, such TNF stimulation likely also triggers NF-κB activation and provides a second wave of transcriptional activity after the initial TCR induced activation. These distinct waves of NF-κB activity may contribute to the apparent heirachy of NF-κB target gene regulation we observed here, especially if TCR and TNF induced NF-κB exhibit distinct requirements and thresholds of IKK activation. This is a possibility given that the ubiquitin scaffolds that form following TCR induced CBM complex formation and TNFR complex I represent highly distinct structures that may vary with the capacity to recruit TAB/TAK and IKK complexes and trigger downstream NF-κB activation. In this context, it may be significant that in vivo, RIPK1 inactivation only partially rescues the T cell response by IKK2 deficient F5 T cells. This suggests that the reduced T cell response is the result of suboptimal NF-κB activation in the absence of IKK2. A1 has been suggested as an important regulator of intrinsic apoptosis in activated T cells(Koenen et al., 2013), so the reduced expression of this NF-κB target may therefore be relevant in this in vivo context. Regardless of mechanism, these data do show that, in addition to regulation of cell death pathways, IKK2 dependent activation of NF-κB is also important for optimal effector expansion in vivo. Identifying the relevant NF-κB targets that mediate this function will be important area of future investigation.

## Materials and methods

### Mice

Mice with the following mutations were used in this study; conditional alleles of *Ikbkb* (Li et al., 2003)*, Chuk* (Gareus et al., 2007), *Tnfaip3* (Vereecke et al., 2010)*, Cre* transgenes expressed under the control of the human CD2 (*huCD2^iCre^*)(de Boer et al., 2003) or *Cd4* expression elements (*Cd4^Cre^*), mice with null mutations for *Tnf*, *Rag1*, *Tnfrsf1*, (Jax Laboratories) mice expressing F5 TCR transgenes(Mamalaki et al., 1993), mice with a D138N mutation in *Ripk1* (RIPK1^D138N^) (Newton et al., 2014). The following strains using combinations of these alllele were bred; F5 *Rag1*^-/-^ *Ikbkb^fx^*^/fx^ huCD2^iCre^ (**F5 *Ikk2*ΔT^CD2^**), F5 *Rag1*^-/-^ *Ikbkb^fx^*^/fx^ huCD2^iCre^ *Tnfrsf1*^-/-^(**F5 *Ikk2*ΔT^CD2^*Tnfrsf1*^-/-^**), F5 *Rag1*^-/-^ *Ikbkb^fx^*^/fx^ huCD2^iCre^ *Ripk1*^D138N^(**F5 *Ikk2*ΔT^CD2^ RIPK1^D138N^**), F5 *Rag1*^-/-^ *Ikbkb^fx^*^/fx^ huCD2^iCre^ *Tnf*^-/-^(**F5 *Ikk2*ΔT^CD2^*Tnf*^-/-^**), *Tnfaip3^fx^*^/fx^ CD4^Cre^ (***Tnfaip3*ΔT^CD4^**), *Ikbkb^fx/fx^* Cd4^Cre^ (***Ikk2*ΔT^CD4^**), *Tnfaip3^fx^*^/fx^ *Ikbkb^fx/fx^* CD4^Cre^ (***Tnfaip3.Ikk2*ΔT^CD4^**), *Tnfaip3^fx^*^/fx^ *Ikbkb^fx/fx^* CD4^Cre^*Ripk1^D138N^* (***Tnfaip3.Ikk2*ΔT^CD4^ RIPK1^D138N^**), *Tnfaip3^fx^*^/fx^ *Chuk^fx/fx^ Ikbkb^fx/fx^* CD4^Cre^ (***Tnfaip3.Ikk1.Ikk2*ΔT^CD4^**), SJL.C57Bl6/J (**CD45.1**). All mice were bred in the Comparative Biology Unit of the Royal Free UCL campus and at Charles River laboratories, Manston, UK. A/NT/60-68 Influenza A virus was innoculated into mice either by intranasal or intraperitoneal routes at a dose of 5HAU/mouse. Animal experiments were performed according to institutional guidelines and Home Office regulations.

### Flow cytometry and electronic gating strategies

Flow cytometric analysis was performed with 2-5 x 10^6^ thymocytes, 1-5 x 10^6^ lymph node or spleen cells. Cell concentrations of thymocytes, lymph node and spleen cells were determined with a Scharf Instruments Casy Counter. Cells were incubated with saturating concentrations of antibodies in 100 μl of Dulbecco’s phosphate-buffered saline (PBS) containing bovine serum albumin (BSA, 0.1%) for 1hour at 4°C followed by two washes in PBS-BSA. Panels used the following mAb: EF450-conjugated antibody against CD25(ThermoFisher Scientific), PE-conjugated antibody against CD127 (ThermoFisher Scientific), BV785-conjugated CD44 antibody (Biolegend), BV650-conjugated antibody against CD4 (Biolegend), BUV395-conjugated antibody against CD8 (BD Biosciences), BUV737-conjugated antibody against CD24 (BD Biosciences), PerCP-cy5.5-conjugated antibody against TCR (Tonbo Biosciences). Cell viability was determined using LIVE/ DEAD cell stain kit (Invitrogen Molecular Probes), following the manufacturer’s protocol. multi-color flow cytometric staining was analyzed on a LSRFortessa (Becton Dickinson) instrument, and data analysis and color compensations were performed with FlowJo V10 software (TreeStar). The following gating strategies were used : Naive peripheral CD4^+^ T cells - CD4^+^ TCR^hi^ CD44^lo^ CD25^lo^, naive peripheral C84^+^ T cells - CD4^+^ TCR^hi^ CD44^lo^ CD25^lo^, memory phenotype CD4^+^ T cells - CD4+TCRhiCD44hiFoxp3–, regulatory T cells - CD4^+^TCR^hi^Foxp3^+^, mature CD4^+^ and CD8^+^ SP thymocytes were identified as TCR^hi^CD4^+^CD8^-^HSA^lo^ and TCR^hi^CD4^-^CD8^+^HSA^lo^ respectively.

### In vitro culture

Thymocytes and LN T cells were cultured at 37°C with 5% CO2 in RPMI-1640 (Gibco, Invitrogen Corporation, CA) supplemented with 10% (v/v) fetal bovine serum (FBS) (Gibco Invitrogen), 0.1% (v/v) 2-mercaptoethanol βME (Sigma Aldrich) and 1% (v/v) penicillin-streptomycin (Gibco Invitrogen) (RPMI-10). Recombinant TNF was supplemented to cultures at 20ng/ml, unless otherwise stated, and was obtained from Peprotech, with PBS used as vehicle. Inhibitors were used at the following concentrations, unless otherwise stated: IKK2 inhibitor BI605906 (IKK2i) (10µM in 0.1% DMSO vehicle), IKK16 (2µM in 0.1% DMSO), Nec1 (10µM in 0.1% DMSO).

### RNAseq analysis

RNA was isolated from single cell suspensions using the RNeasy Mini Kit (Qiagen) according to the manufacturer’s instructions. RNA integrity was confirmed using Agilent’s 2200 Tapestation. Samples were processed using the SMART-Seq v4 Ultra Low Input RNA Kit (Clontech Laboratories, Inc.). Briefly, cDNA libraries were generated using the SMART (Switching Mechanism at 5’ End of RNA Template) technology which produces full-length PCR amplified cDNA starting from 10ng total RNA. The amplified cDNA was checked for integrity and quantity on the Agilent Bioanalyser using the High Sensitivity DNA kit. 150pg of cDNA was then converted to sequencing library using the Nextera XT DNA (Illumina, San Diego, US). This uses a transposon able to fragment and tag the double-stranded cDNA (Tagmentation), followed by a limited PCR reaction (12 cycles). Libraries to be multiplexed in the same run are pooled in equimolar quantities, calculated from Qubit and Tapestation fragment analysis. Samples were sequenced on the NextSeq 500 instrument (Illumina, San Diego, US) using a 43bp paired end run.

Run data were demultiplexed and converted to fastq files using Illumina’s bcl2fastq Conversion Software v2.19. Fastq files are pre-processed to remove adapter contamination and poor quality sequences (trimmomatic v0.36) before being mapped to a suitable reference genome using the spliced aligner STAR (v2.5b). Mapped data is deduplicated using Picard Tools (v2.7.1), in order to remove reads that are the result of PCR amplification, and remaining reads per transcript are counted by FeatureCounts (v1.4.6p5). Normalisation, modelling and differential expression analysis are then carried out using SARTools (v1.3.2), an integrated QC and DESeq2 BioConductor wrapper. After normalization, reads were displayed as fragments per kilobase of exon per million reads (FPKM).

### Immunoblotting and Complex I immunoprecipitation

3 x 10^7^ total thymocytes were used per condition. For complex I immunoprecipitations (IPs), cells were stimulated with 2 mg/ml 3xFLAG-TNF. Cells were washed two times in ice-cold PBS before lysis in 1 ml of NP-40 lysis buffer (10% glycerol, 1% NP-40, 150 mM NaCl, and 10 mM Tris-HCl [pH 8] supplemented with phosphatase and protease inhibitor cocktail tablets [Roche Diagnostics]). The cell lysates were cleared by centrifugation for 15 min at 4^0^C, and the supernatant was then incubated overnight with FLAG M2 affinity gel at 4°C. The next day, the beads were washed three times in PBS buffer. The beads were then either resuspended in 10µl Laemmli buffer to elute the immune complexes or resuspended in 40µl of DUB/lPP buffer (50 mM Tris-HCl [pH 8], 50 mM NaCl, 5 mM DTT, and 1 mM MnCl2) to remove conjugated ubiquitin chains and phosphorylations on RIPK1. Then, either 1.8 mg USP2 or 800 U Lambda protein phosphatase (New England BioLabs) was added as indicated. Reactions were incubated for 30 min at 30 C and subsequently 30 min at 37°C. IPs and total cell extracts were analyzed by NuPage 3-8% Bis-Tris gel (Invitrogen Novex), transferred onto PVDF membrane (Millipore) and immunoblotted with anti-RIPK1 (Cell Signalling Technology). Immunodetection was performed by incubation with horseradish peroxidise-conjugated anti-rabbit (1:5000) (DAKO) and developed by enhanced chemiluminescence (Millipore).

### Statistics

Statistical analysis, line fitting, regression analysis, and figure preparation were performed using Graphpad Prism 8. Column data compared by unpaired Mann-Witney student’s t test. * p<0.01, ** p<0.001.

## Acknowledgements

We thank UCL Comparative Biology Unit staff for assistance with mouse breeding and maintenance. We thank the following for generously sharing of their mouse strains: Prof Manolis Pasparakis for *Chuk* conditional strain, Prof Michael Karin for *Ikbkb* conditional strain, Prof Vishva Dixit for the RIPK1^D138N^ strain, Prof Albert Baldwin for *Rela* conditional strain and Geert van Loo and Rudi Beyaert for conditional *Tnfaip3* strain. The authors declare no competing financial interests. The work in the Seddon lab is supported by the Medical Research Council UK under programme codes MR/P011225/1.

## Non-standard abbreviation list

CBM: CARD11 BCL10 MALT1
CIAP: Cellular inhibitor of apoptosis protein
IKK: inhibitor of kappa B kinase
IKK2i: Inhibitor of kappa B kinase 2
IκB: Inhibitors of kappa B
IL-7R: Interleukin-7 receptor
Nec1: Necrostatin-1
RIPK1: Receptor-interacting protein kinase 1
TAK1: Transforming growth factor-beta activated kinase 1
TNFR1: Tumour necrosis factor receptor 1
TRADD: Tumour necrosis factor receptor 1-associated death domain protein
TRAF: Tumour necrosis factor receptor-associated factor

## Bibliography

Annibaldi, A., and P. Meier. 2018. Checkpoints in TNF-Induced Cell Death: Implications in Inflammation and Cancer. Trends Mol Med 24:49–65.

Blanchett, S., Y. Dondelinger, A. Barbarulo, M.J.M. Bertrand, and B. Seddon. 2022. Phosphorylation of RIPK1 serine 25 mediates IKK dependent control of extrinsic cell death in T cells. Front Immunol 13:1067164.

Bonizzi, G., and M. Karin. 2004. The two NF-kappaB activation pathways and their role in innate and adaptive immunity. Trends Immunol 25:280–288.

Carty, F., S. Layzell, A. Barbarulo, L. Webb, and B. Seddon. 2022. Tonic IKK signalling regulates naive T cell survival in vivo by both repressing RIPK1 dependent extrinsic cell death pathways and independent activation of NFκB. bioRxiv

Chen, X., J. Willette-Brown, X. Wu, Y. Hu, O.M. Howard, Y. Hu, and J.J. Oppenheim. 2015. IKKalpha is required for the homeostasis of regulatory T cells and for the expansion of both regulatory and effector CD4 T cells. FASEB J 29:443–454.

Coornaert, B., M. Baens, K. Heyninck, T. Bekaert, M. Haegman, J. Staal, L. Sun, Z.J. Chen, P. Marynen, and R. Beyaert. 2008. T cell antigen receptor stimulation induces MALT1 paracaspase-mediated cleavage of the NF-kappaB inhibitor A20. Nat Immunol 9:263–271.

de Boer, J., A. Williams, G. Skavdis, N. Harker, M. Coles, M. Tolaini, T. Norton, K. Williams, K. Roderick, A.J. Potocnik, and D. Kioussis. 2003. Transgenic mice with hematopoietic and lymphoid specific expression of Cre. Eur J Immunol 33:314–325.

Dondelinger, Y., M.A. Aguileta, V. Goossens, C. Dubuisson, S. Grootjans, E. Dejardin, P. Vandenabeele, and M.J. Bertrand. 2013. RIPK3 contributes to TNFR1-mediated RIPK1 kinase-dependent apoptosis in conditions of cIAP1/2 depletion or TAK1 kinase inhibition. Cell Death Differ 20:1381–1392.

Dondelinger, Y., S. Jouan-Lanhouet, T. Divert, E. Theatre, J. Bertin, P.J. Gough, P. Giansanti, A.J. Heck, E. Dejardin, P. Vandenabeele, and M.J. Bertrand. 2015. NF-kappaB-Independent Role of IKKalpha/IKKbeta in Preventing RIPK1 Kinase-Dependent Apoptotic and Necroptotic Cell Death during TNF Signaling. Mol Cell 60:63–76.

Dondelinger, Y., P. Vandenabeele, and M.J. Bertrand. 2016. Regulation of RIPK1’s cell death function by phosphorylation. Cell Cycle 15:5–6.

Draber, P., S. Kupka, M. Reichert, H. Draberova, E. Lafont, D. de Miguel, L. Spilgies, S. Surinova, L. Taraborrelli, T. Hartwig, E. Rieser, L. Martino, K. Rittinger, and H. Walczak. 2015. LUBAC-Recruited CYLD and A20 Regulate Gene Activation and Cell Death by Exerting Opposing Effects on Linear Ubiquitin in Signaling Complexes. Cell Rep 13:2258–2272.

Duwel, M., V. Welteke, A. Oeckinghaus, M. Baens, B. Kloo, U. Ferch, B.G. Darnay, J. Ruland, P. Marynen, and D. Krappmann. 2009. A20 negatively regulates T cell receptor signaling to NF-kappaB by cleaving Malt1 ubiquitin chains. J Immunol 182:7718–7728.

Egawa, T., B. Albrecht, B. Favier, M.J. Sunshine, K. Mirchandani, W. O’Brien, M. Thome, and D.R. Littman. 2003. Requirement for CARMA1 in antigen receptor-induced NF-kappa B activation and lymphocyte proliferation. Curr Biol 13:1252–1258.

Fischer, J.C., V. Otten, M. Kober, C. Drees, M. Rosenbaum, M. Schmickl, S. Heidegger, R. Beyaert, G. van Loo, X.C. Li, C. Peschel, M. Schmidt-Supprian, T. Haas, S. Spoerl, and H. Poeck. 2017. A20 Restrains Thymic Regulatory T Cell Development. The Journal of Immunology 199:2356–2365.

Gareus, R., M. Huth, B. Breiden, A. Nenci, N. Rosch, I. Haase, W. Bloch, K. Sandhoff, and M. Pasparakis. 2007. Normal epidermal differentiation but impaired skin-barrier formation upon keratinocyte-restricted IKK1 ablation. Nat Cell Biol 9:461–469.

Gerondakis, S., and U. Siebenlist. 2010. Roles of the NF-kappaB pathway in lymphocyte development and function. Cold Spring Harb Perspect Biol 2:a000182.

Giordano, M., R. Roncagalli, P. Bourdely, L. Chasson, M. Buferne, S. Yamasaki, R. Beyaert, G.v. Loo, N. Auphan-Anezin, A.-M. Schmitt-Verhulst, and G. Verdeil. 2014. The tumor necrosis factor alpha-induced protein 3 (TNFAIP3, A20) imposes a brake on antitumor activity of CD8 T cells. PNAS 111:11115–11120.

Grumont, R., P. Lock, M. Mollinari, F.M. Shannon, A. Moore, and S. Gerondakis. 2004. The Mitogen-Induced Increase in T Cell Size Involves PKC and NFAT Activation of Rel/ NF-κB-Dependent c-myc Expression. Immunity 21:19–30.

Hara, H., T. Wada, C. Bakal, I. Kozieradzki, S. Suzuki, N. Suzuki, M. Nghiem, E.K. Griffiths, C. Krawczyk, B. Bauer, F. D’Acquisto, S. Ghosh, W.C. Yeh, G. Baier, R. Rottapel, and J.M. Penninger. 2003. The MAGUK family protein CARD11 is essential for lymphocyte activation. Immunity 18:763–775.

Hayden, M.S., and S. Ghosh. 2011. NF-kappaB in immunobiology. Cell Res 21:223–244.

Jost, P.J., S. Weiss, U. Ferch, O. Gross, T.W. Mak, C. Peschel, and J. Ruland. 2007. Bcl10/Malt1 signaling is essential for TCR-induced NF-kappaB activation in thymocytes but dispensable for positive or negative selection. J Immunol 178:953–960.

Just, S., G. Nishanth, J.H. Buchbinder, X. Wang, M. Naumann, I. Lavrik, and D. Schluter. 2016. A20 Curtails Primary but Augments Secondary CD8(+) T Cell Responses in Intracellular Bacterial Infection. Sci Rep 6:39796.

Khoshnan, A., C. Tindell, I. Laux, D. Bae, B. Bennett, and A.E. Nel. 2000. The NF-kappa B cascade is important in Bcl-xL expression and for the anti-apoptotic effects of the CD28 receptor in primary human CD4+ lymphocytes. J Immunol 165:1743–1754.

Koenen, P., S. Heinzel, E.M. Carrington, L. Happo, W.S. Alexander, J.G. Zhang, M.J. Herold, C.L. Scott, A.M. Lew, A. Strasser, and P.D. Hodgkin. 2013. Mutually exclusive regulation of T cell survival by IL-7R and antigen receptor-induced signals. Nat Commun 4:1735.

Li, Z.W., S.A. Omori, T. Labuda, M. Karin, and R.C. Rickert. 2003. IKK beta is required for peripheral B cell survival and proliferation. J Immunol 170:4630–4637.

Liu, H.H., M. Xie, M.D. Schneider, and Z.J. Chen. 2006. Essential role of TAK1 in thymocyte development and activation. Proceedings of the National Academy of Sciences of the United States of America 103:11677–11682.

Mamalaki, C., J. Elliott, T. Norton, N. Yannoutsos, A.R. Townsend, P. Chandler, E. Simpson, and D. Kioussis. 1993. Positive and negative selection in transgenic mice expressing a T-cell receptor specific for influenza nucleoprotein and endogenous superantigen. Dev Immunol 3:159–174.

Mora, A.L., R.A. Corn, A.K. Stanic, S. Goenka, M. Aronica, S. Stanley, D.W. Ballard, S. Joyce, and M. Boothby. 2003. Antiapoptotic function of NF-kappaB in T lymphocytes is influenced by their differentiation status: roles of Fas, c-FLIP, and Bcl-xL. Cell Death Differ 10:1032–1044.

Newton, K., D.L. Dugger, K.E. Wickliffe, N. Kapoor, M.C. de Almagro, D. Vucic, L. Komuves, R.E. Ferrando, D.M. French, J. Webster, M. Roose-Girma, S. Warming, and V.M. Dixit. 2014. Activity of protein kinase RIPK3 determines whether cells die by necroptosis or apoptosis. Science 343:1357–1360.

Oh, H., Y. Grinberg-Bleyer, W. Liao, D. Maloney, P. Wang, Z. Wu, J. Wang, D.M. Bhatt, N. Heise, R.M. Schmid, M.S. Hayden, U. Klein, R. Rabadan, and S. Ghosh. 2017. An NF-κB Transcription-Factor-Dependent Lineage-Specific Transcriptional Program Promotes Regulatory T Cell Identity and Function. Immunity 47:450–465.e455.

Onizawa, M., S. Oshima, U. Schulze-Topphoff, J.A. Oses-Prieto, T. Lu, R. Tavares, T. Prodhomme, B. Duong, M.I. Whang, R. Advincula, A. Agelidis, J. Barrera, H. Wu, A. Burlingame, B.A. Malynn, S.S. Zamvil, and A. Ma. 2015. The ubiquitin-modifying enzyme A20 restricts ubiquitination of the kinase RIPK3 and protects cells from necroptosis. Nature Immunology 16:618–627.

Priem, D., M. Devos, S. Druwe, A. Martens, K. Slowicka, A.T. Ting, M. Pasparakis, W. Declercq, P. Vandenabeele, G. van Loo, and M.J.M. Bertrand. 2019. A20 protects cells from TNF-induced apoptosis through linear ubiquitin-dependent and -independent mechanisms. Cell Death Dis 10:692.

Ruefli-Brasse, A.A., D.M. French, and V.M. Dixit. 2003. Regulation of NF-kappaB-dependent lymphocyte activation and development by paracaspase. Science 302:1581–1584.

Ruland, J., G.S. Duncan, A. Elia, I. del Barco Barrantes, L. Nguyen, S. Plyte, D.G. Millar, D. Bouchard, A. Wakeham, P.S. Ohashi, and T.W. Mak. 2001. Bcl10 is a positive regulator of antigen receptor-induced activation of NF-kappaB and neural tube closure. Cell 104:33–42.

Ruland, J., G.S. Duncan, A. Wakeham, and T.W. Mak. 2003. Differential requirement for Malt1 in T and B cell antigen receptor signaling. Immunity 19:749–758.

Saibil, S.D., R.G. Jones, E.K. Deenick, N. Liadis, A.R. Elford, M.G. Vainberg, H. Baerg, J.R. Woodgett, S. Gerondakis, and P.S. Ohashi. 2007. CD4+ and CD8+ T Cell Survival Is Regulated Differentially by Protein Kinase Cθ, c-Rel, and Protein Kinase B. The Journal of Immunology 178:2932–2939.

Schmidt-Supprian, M., G. Courtois, J. Tian, A.J. Coyle, A. Israel, K. Rajewsky, and M. Pasparakis. 2003. Mature T cells depend on signaling through the IKK complex. Immunity 19:377–389.

Schmidt-Supprian, M., J. Tian, E.P. Grant, M. Pasparakis, R. Maehr, H. Ovaa, H.L. Ploegh, A.J. Coyle, and K. Rajewsky. 2004. Differential dependence of CD4+CD25+ regulatory and natural killer-like T cells on signals leading to NF-kappaB activation. Proc Natl Acad Sci U S A 101:4566–4571.

Silva, A., G. Cornish, S.C. Ley, and B. Seddon. 2014. NF-kappaB signaling mediates homeostatic maturation of new T cells. Proc Natl Acad Sci U S A 111:E846–855.

Skaug, B., J. Chen, F. Du, J. He, A. Ma, and Z.J. Chen. 2011. Direct, noncatalytic mechanism of IKK inhibition by A20. Mol Cell 44:559–571.

Sriskantharajah, S., M.P. Belich, S. Papoutsopoulou, J. Janzen, V. Tybulewicz, B. Seddon, and S.C. Ley. 2009. Proteolysis of NF-kappaB1 p105 is essential for T cell antigen receptor-induced proliferation. Nat Immunol 10:38–47.

Ting, A.T., and M.J.M. Bertrand. 2016. More to Life than NF-kappaB in TNFR1 Signaling. Trends Immunol 37:535–545.

Vandenabeele, P., W. Declercq, F. Van Herreweghe, and T. Vanden Berghe. 2010. The role of the kinases RIP1 and RIP3 in TNF-induced necrosis. Sci Signal 3:re4.

Vereecke, L., M. Sze, C. Mc Guire, B. Rogiers, Y. Chu, M. Schmidt-Supprian, M. Pasparakis, R. Beyaert, and G. van Loo. 2010. Enterocyte-specific A20 deficiency sensitizes to tumor necrosis factor-induced toxicity and experimental colitis. J Exp Med 207:1513–1523.

Wan, Y.Y., H. Chi, M. Xie, M.D. Schneider, and R.A. Flavell. 2006. The kinase TAK1 integrates antigen and cytokine receptor signaling for T cell development, survival and function. Nature immunology 7:851–858.

Wang, L., F. Du, and X. Wang. 2008. TNF-alpha induces two distinct caspase-8 activation pathways. Cell 133:693–703.

Webb, L.V., A. Barbarulo, J. Huysentruyt, T. Vanden Berghe, N. Takahashi, S. Ley, P. Vandenabeele, and B. Seddon. 2019. Survival of Single Positive Thymocytes Depends upon Developmental Control of RIPK1 Kinase Signaling by the IKK Complex Independent of NF-κB. Immunity 50:348–361.e344.

Webb, L.V., S.C. Ley, and B. Seddon. 2016. TNF activation of NF-kappaB is essential for development of single-positive thymocytes. J Exp Med 213:1399–1407.

Wertz, I.E., K.M. O’Rourke, H. Zhou, M. Eby, L. Aravind, S. Seshagiri, P. Wu, C. Wiesmann, R. Baker, D.L. Boone, A. Ma, E.V. Koonin, and V.M. Dixit. 2004. De-ubiquitination and ubiquitin ligase domains of A20 downregulate NF-kappaB signalling. Nature 430:694–699.

Xing, Y., X. Wang, S.C. Jameson, and K.A. Hogquist. 2016. Late stages of T cell maturation in the thymus involve NF-κB and tonic type I interferon signaling. Nature Immunology 17:565–573.

Zhang, N., and Y.W. He. 2005. The antiapoptotic protein Bcl-xL is dispensable for the development of effector and memory T lymphocytes. J Immunol 174:6967–6973.

Zheng, Y., M. Vig, J. Lyons, L. Van Parijs, and A.A. Beg. 2003. Combined deficiency of p50 and cRel in CD4+ T cells reveals an essential requirement for nuclear factor kappaB in regulating mature T cell survival and in vivo function. The Journal of Experimental Medicine 197:861–874.

